# Epigenetic gestational age and trajectories of weight and height during childhood: a prospective cohort study

**DOI:** 10.1101/599738

**Authors:** Harold D. Bright, Laura D. Howe, Jasmine N. Khouja, Andrew J. Simpkin, Matthew Suderman, Linda M. O’Keeffe

## Abstract

**Background:** Differences between an individual’s estimated epigenetic gestational age (EGA) and their actual gestational age (GA) are defined as gestational age acceleration (GAA). GAA is associated with increased birthweight and birth length. Whether these associations persist through childhood is yet to be investigated.

**Methods:** We examined the association between GAA and trajectories of height and weight from birth to 10 years (n=785) in a British birth cohort study, the Avon Longitudinal Study of Parents and Children (ALSPAC). EGA of participants was estimated using DNA methylation data from cord blood using a recently-developed prediction model. GA of participants was gathered in ALSPAC from clinical records and was measured from last menstrual period (LMP) for most participants. GAA of participants, measured in weeks, was calculated as the residuals from a regression model of EGA on actual GA. Height and weight were obtained from several sources including birth records, research clinics, routine child health clinics, links to health visitor records and parent-reported measures from questionnaires. Analyses were performed using linear spline multilevel models and adjusted for maternal age, maternal pre-pregnancy BMI, maternal smoking during pregnancy and maternal education.

**Results:** In adjusted analyses, offspring with a one-week greater GAA were born on average 0.14 kg heavier (95% Confidence Interval (CI) 0.09, 0.19) and 0.55 cm taller (95% CI 0.33, 0.78) at birth. These differences in weight persisted up to approximately age 9 months but thereafter began to attenuate and reduce in magnitude. From age 5 years onwards, the association between GAA and weight reversed such that GAA was associated with lower weight and this association strengthened with age (mean difference at age 10 years −0.60 kg (95% CI, −1.19, −0.01)). Differences in height persisted only up to age 9 months (mean difference at 9 months 0.15 cm, (95% CI −0.09, 0.39)). From age 9 months to age 10 years, offspring with a one-week greater GAA were of comparable height to those with no GAA (mean difference at age 10 years −0.07 cm, (95% CI −0.64, 0.50)).

**Conclusions:** Gestational age acceleration is associated with increased birth weight and length and these differences persist to age 9 months. From 5 years onwards, the association of GAA and weight reverses such that by age 10 years greater GAA is associated with lower childhood weight. Further work is required to examine whether the weight effects of GAA strengthen further through adolescence and into early adulthood.

## Introduction

Epigenetic modifications refer to changes that do not affect the underlying DNA base sequence but may affect gene function and phenotypic expression (1). DNA methylation (DNAm) is one type of epigenetic mechanism involving methylation of the cytosine nucleotide at cytosine-phosphate-guanine (CpG) dinucleotide sites (1). Environmental exposures such as tobacco smoking (2) and alcohol consumption (3) are associated with altered patterns of DNAm. DNAm levels also change with age across some CpG sites. DNAm has therefore been used to predict age in both children and adults (4, 5). Differences between an individual’s epigenetic age and their chronological age are defined as age acceleration (AA). Positive AA (i.e. having a higher epigenetic than chronological age) is generally associated with a variety of adverse health outcomes including obesity (6), Alzheimer’s disease (7), cancer (8), lower physical and cognitive fitness (9), and all-cause mortality (9).

Recently, the concept of epigenetic age was extended by Bohlin et al. (10) and Knight et al. (11) using DNAm patterns in cord blood to calculate epigenetic gestational age (EGA), an estimation of gestational age at birth. Differences between an individual’s EGA and their actual gestational age (GA) are defined as gestational age acceleration (GAA). In a recent study of the Avon Longitudinal Study of Parents and Children (ALSPAC), GAA was associated with higher birthweight and greater birth length (12). Given that childhood height and weight are important indicators of overall health and development (13-15), understanding whether associations of GAA with growth measures persist through childhood may provide insights into the potential impact of GAA on later health in children.

In this study we examine the association between GAA and trajectories of weight and height from birth to age 10 years, using data derived from the Accessible Resource for Integrated Epigenomic Studies (ARIES) project, a sub-study of 1,018 mother-offspring pairs from the ALSPAC.

## Methods

### Study Participants

This study uses data collected in ALSPAC, a prospective birth cohort study in southwest England (16, 17). ALSPAC recruited 14,541 pregnant women resident in Avon with expected delivery dates between 1^st^ April 1991 and 31^st^ December 1992. Of these initial pregnancies, there were 14,062 live births and 13,988 children who were alive at 1 year of age. Follow-up has included parent and child completed questionnaires, links to routine data and clinic attendance. Research clinics were held when the participants were approximately 7, 9, 10, 11, 13, 15, and 18 years old. The study has been described elsewhere in detail (16, 17). Ethical approval for the study was obtained from the ALSPAC Ethics and Law Committee and the Local Research Ethics Committees. The study website contains details of all the data that is available through a fully searchable data dictionary and variable search tool (http://www.bristol.ac.uk/alspac/researchers/our-data/). In a sub-study of 1,018 ALSPAC mother-offspring pairs (the ARIES project) (18), genome-wide DNA methylation profiling was performed on offspring DNA samples at birth.

### DNA Methylation

DNA samples at birth were obtained from cord blood, drawn from the umbilical cord upon delivery in accordance with standard procedures. DNA methylation analysis of these samples was performed using the Illumina Infinium HumanMethylation450K BeadChip assay (19). All DNA methylation wet-lab and pre-processing analyses were performed at the University of Bristol as part of the ARIES project. Following extraction, DNA was bisulfite-converted using the Zymo EZ DNA MethylationTM kit (Zymo, Irvine, CA). The Illumina 450K array was used to quantify DNA methylation at over 485,000 CpG sites across the genome. The arrays were then scanned using an Illumina iScan and initial quality review was assessed using GenomeStudio. Samples then underwent a number of further quality control processes. For each sample, the estimated level of DNA methylation at each CpG site was reported as a beta value (β), ranging from 0 (no cytosine methylation) to 1 (complete cytosine methylation) (18).

### Epigenetic Gestational Age and Gestational Age Acceleration

EGA of participants was estimated from DNA methylation data using the Bohlin et al. epigenetic clock model (10). This model uses 96 CpG sites to estimate GA at birth from cord blood DNA methylation. The Bohlin et al. model was used here instead of the Knight et al. model (11) due to its stronger correlation with GA in ARIES (R=0.65 compared to R=0.37) (12, 20). GA of subjects was gathered in ALSPAC from clinical records and was measured from last menstrual period (LMP) for the majority of subjects, although this measure was updated for some individuals following a dating ultrasound. GAA of participants, measured in weeks, was calculated as the residuals from a regression model of EGA on actual GA (12). It is not known which method of GA measurement was used for different individuals; however, the proportion of subjects in which GA was updated from ultrasound is likely to be small as this was not common practice at the time of measurement (12). A positive GAA value corresponds to an EGA which is greater than actual GA. Conversely, a negative GAA value corresponds to an EGA which is less than actual GA.

### Measurement of Height and Weight

Birthweight, birth length, height and weight were collected from several sources from birth to 10 years, including measurement by trained ALPSAC staff shortly after birth, research clinics, routine child health clinics, links to health visitor records and parent-reported measures from questionnaires. Full details of measurement of height and weight are included in Supplementary Material.

### Statistical Analyses

We used multilevel models to examine the association between GAA and trajectories of height and weight from birth to age 10 years. Multilevel models estimate both average and individual-specific trajectories whilst accounting for the change in scale and variance of growth measures over time and the non-independence of repeated measures within individuals (21). Multilevel models also account for differences in the number and timing of measurements between individuals, allowing all available data from eligible participants to be included under a Missing at Random (MAR) assumption (i.e. that the missingness is not specifically related to participants’ height/weight), thereby maximising the number of included participants and minimising selection bias.

Height/weight trajectories were previously modelled (22) and estimated using random-effects linear spline multilevel models (two levels: measurement occasion and individual). Linear spline models divide the growth trajectory into periods in which change is approximately linear (22). With random intercept and random effects for each spline (slope term), individual trajectories are flexible about the mean trend. For length/height growth, knots were placed at 3 months, 12 months and 36 months, resulting in four periods of change; birth to 3 months, 3 to 12 months, 12 to 36 months and 36 to 120 months. For weight change, an additional knot was placed at 84 months, resulting in five periods of change; birth to 3 months, 3 to 12 months, 12 to 36 months, 36 to 84 months and 84 to 120 months. Trajectories were modelled separately for males and females, and only up to age 10 years due to variability in age at puberty onset which would require knot points to be placed at different ages for each individual. Further details are included in Supplementary Material.

The associations between GAA and the height/weight trajectories were modelled by including interaction terms between GAA and the intercept (birth length/weight) and slopes (rate of height/weight growth in each linear spline period). We performed unadjusted and confounder-adjusted analyses. We considered maternal pre-pregnancy BMI, maternal age, maternal education and maternal smoking during pregnancy as potential confounders (all measured by mother-completed questionnaires; details in Supplementary Material). Covariates were included as interactions between each covariate and the intercept and linear slopes in the multilevel models. All participants with a measure of GAA, at least one measure of height/weight from birth to 10 years and complete data on all covariates were included, leading to a total sample of 785 (398 females and 387 males) included in analyses. All analyses were conducted using the statistical packages Stata version 15.1 (23), MLwiN version 3.01 (24) and the Stata command ‘runmlwin’ (25).

### Sensitivity Analyses

We performed sensitivity analyses, examining the association of GAA with observed measures of height/weight at birth, age 3 years and age 10 years using linear regression to examine the suitability of our modelling strategy. We also compared characteristics of participants who were included in our analysis versus participants who were excluded due to missing data on exposure, outcome or confounders.

## Results

### Study Participants

Characteristics of participants included in our analysis are shown in Table 1. Mothers of offspring included in the analysis (n=785) were likely to be older, be married, have attained a higher level of education and were less to likely to smoke during pregnancy relative to the rest of the ALSPAC study population excluded due to missing data on exposure, outcome or confounders (n=13,446) (Supplementary Table 2). Included participants had a total of 9,515 height and weight measurements with a median of 13 measures per individual (interquartile range (IQR) 9 to 19).

**Table 1.** Characteristics of the study sample (n=785).

|  | n | % of study sample |
| --- | --- | --- |
| **Sex of Child** |  |  |
| Male | 387 | 49.3 |
| Female | 398 | 50.7 |
| **Maternal age (years)** |  |  |
| 35+ | 107 | 13.6 |
| 25-34 | 586 | 74.7 |
| 15-24 | 92 | 11.7 |
| **Maternal education\*** |  |  |
| Degree or above | 160 | 20.4 |
| A level | 242 | 30.8 |
| O level | 258 | 32.9 |
| Less than O level | 125 | 15.9 |
| **Maternal smoking during pregnancy** |  |  |
| Never | 477 | 60.8 |
| Ever | 308 | 39.2 |
| **Maternal pre-pregnancy BMI** |  |  |
| Under/normal weight | 637 | 81.2 |
| Overweight | 111 | 14.1 |
| Obese | 37 | 4.7 |
| **Maternal marital status during pregnancy\*\*** |  |  |
| Never married | 93 | 11.9 |
| Married | 650 | 83.1 |
| Divorced/separated/widowed | 39 | 5.0 |
| **Parity\*\*** |  |  |
| 0 | 651 | 84.3 |
| 1 | 93 | 12.1 |
| 2+ | 28 | 3.6 |
|  | **Median** | **IQR\*\*\*** |
| **Gestational age at birth (weeks)** | 40 | 39, 41 |

\* O-Level (Ordinary Level; exams taken in different subjects usually at age 15–16 at the completion of legally required school attendance, equivalent to today's UK General Certificate of Secondary Education), A-Level (Advanced-Level; exams taken in different subjects usually at age 18), or university degree or above.
\*\* Denominator less than 785 as data not required for inclusion in the study population.
\*\*\* Interquartile range.

**Table 2.**
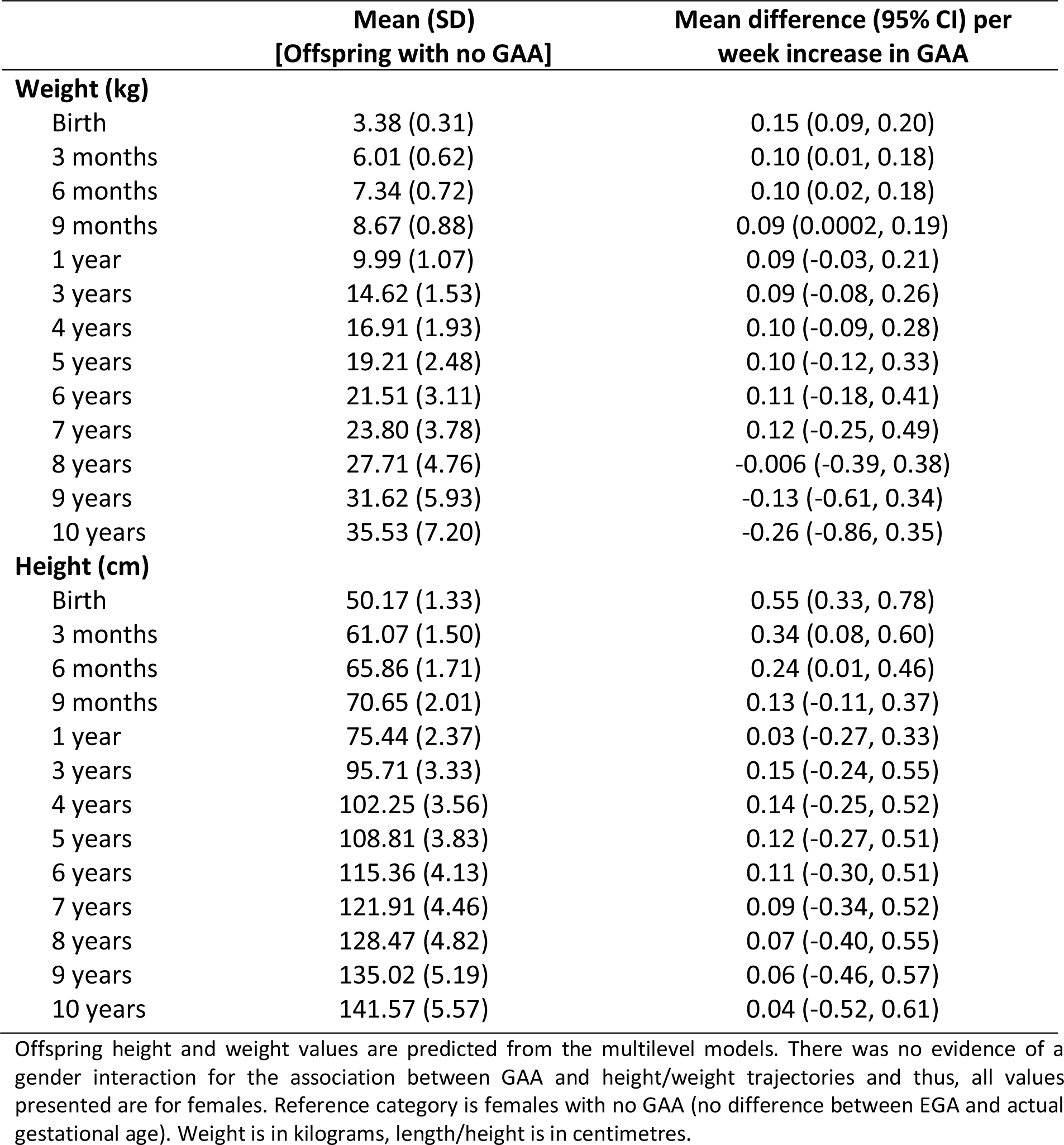
Unadjusted association of gestational age acceleration (GAA) with predicted offspring weight and length/height at multiple ages (n=785).

|  | Mean (SD)<br>[Offspring with no GAA] | Mean difference (95% CI) per<br>week increase in GAA |
| --- | --- | --- |
| **Weight (kg)** |  |  |
| Birth | 3.38 (0.31) | 0.15 (0.09, 0.20) |
| 3 months | 6.01 (0.62) | 0.10 (0.01, 0.18) |
| 6 months | 7.34 (0.72) | 0.10 (0.02, 0.18) |
| 9 months | 8.67 (0.88) | 0.09 (0.0002, 0.19) |
| 1 year | 9.99 (1.07) | 0.09 (-0.03, 0.21) |
| 3 years | 14.62 (1.53) | 0.09 (-0.08, 0.26) |
| 4 years | 16.91 (1.93) | 0.10 (-0.09, 0.28) |
| 5 years | 19.21 (2.48) | 0.10 (-0.12, 0.33) |
| 6 years | 21.51 (3.11) | 0.11 (-0.18, 0.41) |
| 7 years | 23.80 (3.78) | 0.12 (-0.25, 0.49) |
| 8 years | 27.71 (4.76) | -0.006 (-0.39, 0.38) |
| 9 years | 31.62 (5.93) | -0.13 (-0.61, 0.34) |
| 10 years | 35.53 (7.20) | -0.26 (-0.86, 0.35) |
| **Height (cm)** |  |  |
| Birth | 50.17 (1.33) | 0.55 (0.33, 0.78) |
| 3 months | 61.07 (1.50) | 0.34 (0.08, 0.60) |
| 6 months | 65.86 (1.71) | 0.24 (0.01, 0.46) |
| 9 months | 70.65 (2.01) | 0.13 (-0.11, 0.37) |
| 1 year | 75.44 (2.37) | 0.03 (-0.27, 0.33) |
| 3 years | 95.71 (3.33) | 0.15 (-0.24, 0.55) |
| 4 years | 102.25 (3.56) | 0.14 (-0.25, 0.52) |
| 5 years | 108.81 (3.83) | 0.12 (-0.27, 0.51) |
| 6 years | 115.36 (4.13) | 0.11 (-0.30, 0.51) |
| 7 years | 121.91 (4.46) | 0.09 (-0.34, 0.52) |
| 8 years | 128.47 (4.82) | 0.07 (-0.40, 0.55) |
| 9 years | 135.02 (5.19) | 0.06 (-0.46, 0.57) |
| 10 years | 141.57 (5.57) | 0.04 (-0.52, 0.61) |

Offspring height and weight values are predicted from the multilevel models. There was no evidence of a gender interaction for the association between GAA and height/weight trajectories and thus, all values presented are for females. Reference category is females with no GAA (no difference between EGA and actual gestational age). Weight is in kilograms, length/height is in centimetres.

### Weight

In the unadjusted analysis (Table 2), a one-week greater GAA was associated with 0.15 kg higher birth weight (95% confidence interval (CI) 0.09, 0.20). Offspring with a one-week greater GAA continued to be heavier than those with no GAA up to 7 years (mean difference 0.12 kg, 95% CI −0.25, 0.49), albeit with confidence intervals that spanned the null value from age 1 year onwards. From age 8 years, a one-week greater GAA was associated with lower weight although this difference spanned the null value throughout (mean difference at age 10 −0.26 kg, 95% CI −0.86, 0.35). In the confounder-adjusted analysis (Table 3), associations were similar but with greater evidence for a reversal of the association of GAA and weight from positive to negative across childhood. Weight differences for a one-week greater GAA were present at birth, persisted up to age up to 4 years (mean difference 0.02 kg, 95% CI −0.16, 0.21) but with confidence intervals that spanned the null from 9 months. From age 5 years, the association reversed such that GAA was associated with lower weight and this association strengthened with age (mean difference at age 10 −0.60 kg, 95% CI − 1.19, −0.01). A one-week greater GAA was associated with a slower rate of weight growth from birth onwards, albeit with confidence intervals that spanned the null (Supplementary Table 4). This explains the attenuation of differences in weight by GAA at age 9 months and eventual reversal of the association between GAA and weight at age 5 years.

**Table 3.** Confounder-adjusted association of gestational age acceleration (GAA) with predicted offspring weight and length/height at multiple ages (n=785).

|  | Mean (SD)<br>[Offspring with no GAA] | Mean difference (95% CI) per<br>week increase in GAA |
| --- | --- | --- |
| **Weight (kg)** |  |  |
| Birth | 3.44 (0.30) | 0.14 (0.09, 0.19) |
| 3 months | 5.98 (0.61) | 0.10 (0.02, 0.19) |
| 6 months | 7.29 (0.71) | 0.09 (0.01, 0.18) |
| 9 months | 8.61 (0.87) | 0.09 (-0.01, 0.18) |
| 1 year | 9.92 (1.06) | 0.08 (-0.04, 0.20) |
| 3 years | 14.44 (1.50) | 0.05 (-0.12, 0.23) |
| 4 years | 16.70 (1.87) | 0.02 (-0.16, 0.21) |
| 5 years | 18.95 (2.39) | -0.008 (-0.23, 0.22) |
| 6 years | 21.21 (2.98) | -0.04 (-0.33, 0.25) |
| 7 years | 23.47 (3.61) | -0.07 (-0.43, 0.29) |
| 8 years | 27.26 (4.52) | -0.25 (-0.63, 0.13) |
| 9 years | 31.05 (5.63) | -0.43 (-0.89, 0.04) |
| 10 years | 34.85 (6.83) | -0.60 (-1.19, -0.01) |
| **Height (cm)** |  |  |
| Birth | 50.41 (1.34) | 0.55 (0.33, 0.78) |
| 3 months | 60.96 (1.50) | 0.35 (0.09, 0.62) |
| 6 months | 65.69 (1.71) | 0.25 (0.02, 0.48) |
| 9 months | 70.41 (2.01) | 0.15 (-0.09, 0.39) |
| 1 year | 75.13 (2.37) | 0.05 (-0.26, 0.35) |
| 3 years | 95.24 (3.29) | 0.11 (-0.28, 0.50) |
| 4 years | 101.78 (3.51) | 0.09 (-0.30, 0.47) |
| 5 years | 108.32 (3.78) | 0.06 (-0.33, 0.45) |
| 6 years | 114.85 (4.08) | 0.03 (-0.37, 0.44) |
| 7 years | 121.39 (4.41) | 0.008 (-0.43, 0.44) |
| 8 years | 127.92 (4.76) | -0.02 (-0.49, 0.46) |
| 9 years | 134.46 (5.13) | -0.04 (-0.56, 0.47) |
| 10 years | 140.99 (5.51) | -0.07 (-0.64, 0.50) |

Offspring height and weight values are predicted from the multilevel models, based on models adjusted for maternal age, maternal smoking during pregnancy, maternal education and maternal pre-pregnancy BMI. There was no evidence of a gender interaction for the association between GAA and height/weight trajectories and thus, all values presented are for females. Reference category is females with no GAA (no difference between EGA and actual gestational age). Weight is in kilograms, length/height is in centimetres.

### Length/Height

In the unadjusted analysis (Table 2), a one-week greater GAA was associated with 0.55 cm greater birth length (95% CI 0.33, 0.78). Differences in height for a one-week greater GAA diminished with age and confidence intervals spanned the null value from age 9 months onward. In the confounder-adjusted analysis (Table 3), findings were similar but there was some evidence of a reversal of the association of GAA and height from age 8 years, albeit with confidence intervals that spanned the null value (mean difference at age 10 −0.07 cm, 95% CI −0.64, 0.50). A one-week greater GAA was associated with a slower rate of height growth from birth onwards, albeit with confidence intervals that spanned the null (Supplementary Table 4). This explains the attenuation of differences in height by GAA at age 9 months and eventual reversal of the association between GAA and height from age 8 years.

### Sensitivity Analysis

Examination of the association of GAA with observed measures of weight and height (rather than predicted measures from multilevel models) at birth, age 3 years and age 10 years using linear regression produced similar results, providing reassurance that our modelling strategy was appropriate (Supplementary Table 3).

## Discussion

### Summary

Using cord blood DNAm at birth to calculate EGA, we examined the association between GAA and trajectories of childhood weight and height from birth to 10 years in the ARIES subsample of ALSPAC. Our findings showed that GAA was associated with higher birthweight and birth length as demonstrated previously (12) and that this difference persisted up to approximately 9 months of age. From age 9 months onwards, this difference continued to attenuate and eventually reversed for weight, resulting in approximately 0.6 kg lower weight at age 10 years per week greater GAA.

### Previous Research

To our knowledge this is the first study to examine the association of GAA with measures of weight and height beyond birth and across childhood. A previous study examining DNA methylation patterns in cord blood DNA found that DNA methylation levels in only 1 of their studied genes was associated with body size at age 9 years (26); a finding which contrasts slightly with the association of GAA and weight from age 5 years found here which strengthened with age. A study of AA also conducted in the ARIES subsample of ALSPAC found that AA at age 7 years was positively associated with height, but not with weight, from 7-17 years (27). The same study found that AA at birth was associated with a higher fat mass from birth to age 17 years but found no evidence of an association between AA and birthweight or length, in contrast to the association between GAA and birthweight/length. However, it should be noted that epigenetic age/AA are uncorrelated with gestational age at delivery (28) and the CpG sites used to measure GAA (10) differ substantially from those used in the Horvath model (4) to measure AA. These findings indicate that AA and GAA are largely independent constructs, offering an explanation for the disparity seen in associations of GAA and AA with birth/childhood anthropometric outcomes.

### Potential implications

Our results indicate that GAA is not strongly associated with increased childhood weight and height for most of childhood and that by age 10 years GAA is associated with lower weight. The attenuation of the association between GAA and body size after birth could be due to the possible reversibility of epigenetic changes i.e. methylation patterns are not fixed and may be modified. This may also support the reversal of the association seen for weight from age 5 years onward. The process of reversibility has been observed previously with the effect of maternal smoking during pregnancy on offspring DNA methylation levels, which were found to have reverted at many CpG sites by age 7 years (29). Further work is required to examine whether the association between GAA and lower weight at age 10 continues to strengthen with age.

The Bohlin et al. model epigenetic clock model measures DNA methylation levels in cord blood only and therefore does not examine DNA methylation changes in other tissues at birth. As tissue-specific age acceleration associations have been observed previously, such as the association between AA in liver cells and obesity (6), investigation of gestational age-related DNA methylation patterns in foetal tissues other than cord blood may also provide important insights.

### Strengths and Limitations

There are several strengths to our study including the use of repeated measures of height and weight from birth to 10 years, and the use of multilevel models allowing all individuals with at least one measure of height and weight to be included in analyses, thereby reducing the possibility of selection bias. The Bohlin et al. (10) epigenetic clock model transferred well to the ARIES study population, achieving a high correlation between EGA and actual GA (R=0.65) (20). Limitations include estimation of GA from LMP rather than the ‘gold standard’ of dating ultrasound, which may have implications for our estimation of EGA, as the Bohlin et al. model produces more accurate EGA estimates in cohorts in which GA has been measured from ultrasound (10). Height and weight data in ALSPAC were obtained from a range of sources with varying degrees of reliability; parent-reported measures from questionnaires will likely have less accuracy than research clinic measurements following strict methodology. However, our models have aimed to minimise this by accounting for differential measurement error across measurement sources (see Supplementary Material for handling of clinic and parent-reported measurements). Participants included in our analysis were more advantaged than those excluded due to missing exposure, outcome or confounder data, introducing the possibility of selection bias and reducing generalisability of findings. The Bohlin et al. model was developed using the HumanMethylation450K BeadChip assay (19) which has some recognised technical biases and only profiles 1.7% of CpG sites in the genome. The assay may therefore have failed to identify some CpGs that are differentially methylated in newborns at birth. The model also applies solely to DNAm in cord blood, which may not be the optimal tissue for relating EGA and physical development.

### Conclusion

Gestational age acceleration is associated with increased birthweight and length with differences persisting to age 9 months. From age 5 years onwards, the association of GAA and weight reverses such that by age 10 years greater GAA is associated with lower weight. Further work is required to examine whether the weight effects of GAA strengthen through adolescence and into early adulthood.

## Declarations

### Ethics approval and consent to participate

Ethical approval for the study was obtained from the ALSPAC Ethics and Law Committee and the Local Research Ethics Committees. Informed consent was given by the offsprings’ parents at the outset of the study.

### Consent for publication

Not applicable.

### Availability of data and materials

The data sets generated and/or analysed during the current study are available from the corresponding author on reasonable request, subject to the ALSPAC study executive data access procedures, as specified on the ALSPAC website (http://www.bristol.ac.uk/alspac/researchers/access/) for researchers who meet the criteria for access to confidential data.

## Authors’ contributions

The study was conceived and designed by LMOK. Statistical analyses were conducted by HDB and LMOK. HDB was responsible for writing the first draft of the manuscript. HDB, LMOK, LDH, JNK, AJS and MS contributed to the interpretation of results, critical revisions of the manuscript and approved the final manuscript.

## Acknowledgements

We are extremely grateful to all the families who took part in this study, the midwives for their help in recruiting them, and the whole ALSPAC team, which includes interviewers, computer and laboratory technicians, clerical workers, research scientists, volunteers, managers, receptionists and nurses.

## Abbreviations

AA: Age acceleration;
ALSPAC: Avon Longitudinal Study of Parents and Children;
ARIES: Accessible Resource for Integrated Epigenomic Studies;
BMI: Body mass index;
CI: Confidence interval;
DNAm: DNA methylation;
EGA: Epigenetic gestational age;
GA: Gestational age;
GAA: Gestational age acceleration;
IQR: Interquartile range;
LMP: Last menstrual period;
SD: Standard deviation;
SE: Standard error.

