## Supplementary Material for "Epigenetic gestational age and trajectories of weight and height during childhood: a prospective cohort study"

**Details of measurement of height and weight at research clinics**

Birth length and weight were available for most children. Birth length (crown-heel) was measured by Avon Longitudinal Study of Parents and Children (ALSPAC) staff soon after birth (median 1 day, range 1-14 days) using a Harpenden Neonatometer (Holtain Ltd). Birthweight was extracted from medical records. Between birth and age 5 years, measures were available from routine child health clinics for most children, and from research clinic measurements on a random 10% subsample of the cohort. All cohort members were invited to research clinics from age 7 onwards. Across all ages parent-reported measures were available from questionnaires. Length and weight measurements were extracted from health visitor records, which form part of standard child care in the UK. At the clinics between 4 months and 5 years, crown-heel length for children aged 4 to 25 months was measured using a Harpenden Neonatometer and from 25 months onwards standing height was measured using a Leicester Height Measure; weight was measured using Fereday 100kg combined scale (4-month clinic), Soenhle scale or Seca scale model 724 (8-month clinic), Seca 724 or Seca 835 (12-month clinic), Seca 835 (18 months onwards). At annual clinics from age 7 onwards, standing height was measured to the last complete mm using the Harpenden Stadiometer and weight was measured to the nearest 0.1kg using the Tanita Body Fat Analyser (Model TBF 305).

**Details of measurement of confounding variables**

A questionnaire at 32 weeks gestation asked mothers to report their educational attainment, which was categorised as below O-Level (Ordinary Level; exams taken in different subjects usually at age 15-16 at the completion of legally required school attendance, equivalent to today’s UK General Certificate of Secondary Education), O-Level only, A-Level (Advanced Level; exams taken in different subjects usually at age 18), or university degree or above. Smoking data were self-reported by mothers during questionnaires at 18 weeks gestation; responses to having ever smoked, as well as having stopped now, were used to categorise maternal smoking status as never smoked or ever smoked during pregnancy. Maternal height and pre-pregnancy weight were self-reported from a questionnaire administered at 12 weeks gestation; these were used to calculate maternal pre-pregnancy BMI. Maternal age was reported in the mother’s antenatal questionnaires.

**Details of previously published work on model selection**

Trajectories of height and weight have been modelled previously and are described elsewhere in detail (1-3). In brief, the standard way of modelling the relationship between a continuous outcome (in our example height and weight) and a continuous exposure (in our example age) would be to fit a polynomial curve (i.e. age raised to the appropriate power). The patterns of growth across childhood, however, follow a complex pattern and may not be accurately represented by a simple polynomial curve. For this reason, we used fractional polynomials to find the best-fitting average height and weight trajectory for each of our models from birth to ten years. Fractional polynomials provide greater modelling flexibility than polynomials by allowing non-integer powers and multiples of logarithms. In our example, age was raised to various combinations of powers (each of the following single powers, plus each combination of two powers: 0.5, 1, 2, 3, -0.5, -1, -2, natural log), from which we selected the best fitting curve (the one with the lowest likelihood value). Although fractional polynomials provide a flexible way to examine such relationships, they do not provide parameters that are clinically relevant, easily interpreted, or readily used to assess the impact of different growth periods on later outcomes. We therefore used the best-fitting fractional polynomial to derive a piecewise linear spline model. From the best fitting curves, we derived approximate knot-points for the linear spline model, i.e. the turning points of the curve between which changes in height and weight were approximately linear.

To optimise the knot points, we fitted a series of models with the knot points placed at 1-month intervals around the estimated knot point identified from the shape of the fractional polynomial curve. The model with the lowest likelihood value was selected as the knot point for the final model. This confirmed that there were:

- four periods of growth (length/height): birth to 3 months, 3 to 12 months, 12 to 36 months, 36 to 120 months.
- five periods of weight change: birth to three months, 3 to 12 months, 12 to 36 months, 36 to 84 months, 84 to 120 months.

To account for the likely reduced accuracy of parent-reported measurements (4), a binary indicator of measurement source (research clinic/health records versus parent-report) was included in the multilevel models. An age term was also included as a random effect at the occasion level, which allowed the measurement error to vary with age.

**Supplementary Table 1.** Assessment of multilevel model fit. Comparison of predicted values (from multilevel models) with observed measures at birth, age 3 years and age 10 years.

|  | Mean observed  (SD) | Mean predicted  (SD) | Mean difference (observed – predicted) | 95% level of agreement between observed and predicted |
| --- | --- | --- | --- | --- |
| Weight (kg) |  |  |  |  |
| Birth | 3.48 (0.48) | 3.38 (0.33) | 0.11 | -0.33, 0.54 |
| 3 years^*^ | 15.31 (1.82) | 14.91 (1.68) | 0.40 | -0.76, 1.56 |
| 10 years^**^ | 34.18 (6.22) | 34.32 (5.85) | -0.14 | -1.66, 1.38 |
| Height (cm) |  |  |  |  |
| Birth | 50.79 (2.04) | 50.11 (1.39) | 0.68 | -1.60, 2.96 |
| 3 years^*^ | 95.95 (3.67) | 95.93 (3.60) | 0.02 | -2.72, 2.76 |
| 10 years^**^ | 140.15 (5.74) | 140.67 (5.29) | -0.52 | -2.87, 1.83 |

* Observed height/weight measures at age 3 years refer to measurements taken from 34-38 months.

** Observed height/weight measures at age 10 years refer to measurements taken from 118-120 months.

**Supplementary Table 2.** Comparison of participants included in the analysis compared with those excluded (i.e. the rest of the ALSPAC cohort) due to missing data on exposure, outcome or confounder.

|  | Participants included  n = 785 | Participants excluded  n = 13,446 | P value for comparison^**^ |
| --- | --- | --- | --- |
|  | n^*^ (%) | n^*^ (%) |  |
| Sex of child |  |  |  |
| Male | 387 (49.30) | 6934 (51.57) | 0.22 |
| Female | 398 (50.70) | 6512 (48.43) |  |
| Maternal age (years) |  |  |  |
| 35+ | 107 (13.63) | 1756 (13.06) | <0.001 |
| 25-34 | 586 (74.65) | 8485 (63.10) |  |
| 15-24 | 92 (11.72) | 3205 (23.84) |  |
| Maternal education^***^ |  |  |  |
| Degree or above | 160 (20.38) | 1413 (12.36) | <0.001 |
| A level | 242 (30.83) | 2494 (21.82) |  |
| O level | 258 (32.87) | 3977 (34.80) |  |
| Less than O level | 125 (15.92) | 3545 (31.02) |  |
| Maternal smoking during pregnancy |  |  |  |
| Never | 477 (60.76) | 5843 (48.51) | <0.001 |
| Ever | 308 (39.24) | 6201 (51.49) |  |
| Maternal pre-pregnancy BMI |  |  |  |
| Under/normal weight | 637 (81.15) | 8350 (79.06) | 0.35 |
| Overweight | 111 (14.14) | 1622 (15.36) |  |
| Obese | 37 (4.71) | 590 (5.59) |  |
| Maternal marital status during pregnancy |  |  |  |
| Never married | 93 (11.89) | 2361 (19.54) | <0.001 |
| Married | 650 (83.12) | 8994 (74.43) |  |
| Divorced/separated/widowed | 39 (4.99) | 729 (6.03) |  |
| Parity |  |  |  |
| 0 | 651 (84.33) | 9490 (79.53) | 0.002 |
| 1 | 93 (12.05) | 1720 (14.41) |  |
| 2+ | 28 (3.63) | 723 (6.06) |  |
|  | **Median (IQR)** | **Median (IQR)** |  |
| Gestational age at birth (weeks) | 40 (39,41) | 40 (39,41) | 0.04 |

* Denominator varies due to missing data.

** P value is for the difference in proportions for categorical variables from Chi^2^ test, or difference in means for continuous variables from t-test between included and excluded participants.

*** O-Level (Ordinary Level; exams taken in different subjects usually at age 15–16 at the completion of legally required school attendance, equivalent to today’s UK General Certificate of Secondary Education), A-Level (Advanced Level; exams taken in different subjects usually at age 18), or university degree or above.

**Supplementary Table 3.** Association of gestational age acceleration with observed weight and length/height at birth, age 3 years and age 10 years using linear regression.

|  | Number of observed measures | Mean (SE)  [Offspring with no GAA] | Mean difference (95% CI) per week increase in GAA |
| --- | --- | --- | --- |
| Weight (kg) |  |  |  |
| Birth | 661 | 3.48 (0.02) | 0.15 (0.10, 0.20) |
| 3 years^*^ | 248 | 15.31 (0.12) | 0.005 (-0.33, 0.34) |
| 10 years^**^ | 227 | 34.20 (0.41) | -0.60 (-1.64, 0.44) |
| Height (cm) |  |  |  |
| Birth | 661 | 50.79 (0.08) | 0.53 (0.32, 0.75) |
| 3 years^*^ | 248 | 95.95 (0.23) | 0.11 (-0.56, 0.78) |
| 10 years^**^ | 227 | 140.15 (0.38) | -0.04 (-1.00, 0.93) |

* Observed height/weight measures at age 3 years refer to measurements taken from 34-38 months.

** Observed height/weight measures at age 10 years refer to measurements taken from 118 -120 months.

**Supplementary Table 4.** Confounder-adjusted association of gestational age acceleration with weight growth and length/height growth from birth to 10 years (n=785).

|  | Mean (SE)  [Offspring with no GAA] | Mean difference (95% CI) per week increase in GAA |
| --- | --- | --- |
| Weight |  |  |
| Birthweight (kg) | 3.44 (0.05) | 0.14 (0.09, 0.19) |
| 0 to 3 months (kg/month) | 0.85 (0.03) | -0.01 (-0.04, 0.02) |
| 3 to 12 months | 0.44 (0.01) | -0.003 (-0.02, 0.01) |
| 12 to 36 months | 0.19 (0.01) | -0.001 (-0.01, 0.01) |
| 36 to 84 months | 0.19 (0.01) | -0.003 (-0.01, 0.005) |
| 84 to 120 months | 0.32 (0.01) | -0.01 (-0.03, -0.001) |
| Height |  |  |
| Birth length (cm) | 50.41 (0.22) | 0.55 (0.33, 0.78) |
| 0 to 3 months (cm/month) | 3.52 (0.09) | -0.07 (-0.16, 0.03) |
| 3 to 12 months | 1.57 (0.03) | -0.03 (-0.07, 0.003) |
| 12 to 36 months | 0.84 (0.01) | 0.003 (-0.01, 0.02) |
| 36 to 120 months | 0.54 (0.01) | -0.002 (-0.01, 0.004) |

Offspring height growth and weight growth are predicted from the multilevel models, based on models adjusted for maternal age, maternal smoking during pregnancy, maternal education and maternal pre-pregnancy BMI. There was no evidence of a gender interaction for the association between GAA and height/weight trajectories and thus, all values presented are for females. Reference category is females with no GAA (no difference between EGA and actual gestational age). Weight is in kilograms, length/height is in centimetres. Weight growth is in kg/month, length/height growth is in cm/month.
